# The Cost of Adaptability: Resource Availability Constrains Functional Stability Under Pulsed Disturbances

**DOI:** 10.1101/2022.12.22.521578

**Authors:** Angel Rain-Franco, Hannes Peter, Guilherme P. de Moraes, Sara Beier

## Abstract

Global change exposes ecosystems to changes in the frequency, magnitude and concomitancy of disturbances, which impact the composition and functioning. Here we experimentally evaluate effects of salinity disturbances and eutrophication on bacterial communities from coastal ecosystems. The resistance, resilience and functional stability of these communities is critically important for water quality, productivity and consequently ecosystem services, such as fishery yields. Yet, little is known about the underlying traits. Microbial functional stability can be maintained via resistance and resilience, which are reflected in genomic traits such as genome size and codon usage bias and may be linked to metabolic costs. To study the impact of pulsed disturbances on community assembly and functioning in dependence of metabolic costs, we performed a 41-days pulse disturbance experiment crossed with two levels of resource availability. Our setup triggered stochastic community re-assembly processes in all treatments. In contrast, we observed consistent and resource availability dependent patterns of superordinate community structural patterns and functioning, such as genomic trait distributions, species diversity, and functional resistance in response to disturbances. Genomic traits reflected the selection for taxa possessing resistant- and resilience-related traits, particularly under high nutrient availability. Our findings thereby mark an important step towards unraveling the compositional and genomic underpinnings of functional resistance in microbial communities after exposure to consecutive pulse disturbances. Our work demonstrates how resource availability alleviates metabolic constraints on resistance and resilience. This has important consequences for predicting water quality and ecosystem productivity of environments exposed to global change.

**Significance:** Understanding the responses of communities to disturbances is a prerequisite to predict ecosystem dynamics and thus highly relevant in light of global change. Microbial communities play key roles in numerous ecosystem functions and services, and the large diversity, rapid growth and phenotypic plasticity of microorganisms are thought to allow for high resistance and resilience. While potential metabolic costs associated with adaptions to fluctuating environments have been debated, little evidence supports trade-offs between resource availability and resistance and resilience. Here, we experimentally assessed the compositional and functional responses of an aquatic microbial model community to disturbances and systematically manipulated resource availability. Our results demonstrate that the capacity to tolerate environmental fluctuations is encoded in genomic traits and constrained by resource availability.

## INTRODUCTION

There is mounting evidence that anthropogenic global change rapidly alters natural disturbance regimes with important consequences for ecosystems and the services they provide (1–3). Particularly, climate warming-induced droughts and heavy precipitation events affect both terrestrial and aquatic ecosystems (4–6). Concomitantly, land-use changes, such as intensified freshwater use for irrigation or eutrophication linked to fertilizer application deteriorate water quality and raise concerns for critical ecosystem services (7–9). Coastal areas are particularly vulnerable to such combined perturbations (7). While interactions between isolated disturbance events, biodiversity and ecosystem consequences have been elucidated (e.g. 10, 11), the underlying traits that allow populations to resist or recover from multiple and consecutive perturbations remain less studied. Although ecological theory exists on the effects of consecutive perturbations (1), this has been developed mainly for larger organisms with long generation times and complex life-histories, while we currently lack an understanding of the effects of frequent disturbances on microbial communities. This is in stark contrast with the important role these microbial communities play for water quality and ecosystem productivity, notably in coastal ecosystems 12/22/22 2:24:00 PM.

Disturbance ecology and biodiversity-ecosystem-functioning (BEF) research has highlighted the importance of species diversity for ensuring functional resistance and resilience (10, 11). The response of microbial communities to disturbances is thought to be determined by the degree to which a community is insensitive to a disturbance (i.e. resistance), and the rate at which a community returns to the pre-disturbance state (13). While functionally redundant taxa can contribute to the resistance and resilience of microbial communities (i.e. insurance effect of diversity), response traits allow individual populations to adapt to and recover from perturbations (14, 15). Of particular interest are life-history traits associated with stress tolerance and adaptation (i.e. resistance traits) as well as traits related to growth and reproduction (i.e. resilience traits)(16).

In line with these expectations, previous experimental work has linked functional stability of microbial communities to the presence of generalist taxa with broad niches (17). It has been suggested that traits inferred from genomic information of microbial communities approximate life history relevant for resistance and resilience (18). For instance, genomic traits such as genome size and the fraction of transcription factors (%TF) were linked to classifications along the generalist-specialist continuum because additional auxiliary genes and regulatory capacity enhance aptness to environmental change (19, 20). In contrast, codon usage bias or the number of 16S rRNA gene copies (RRN) are inversely related to generation time and lag-phase duration (21, 22). Both, short generation times (i.e. rapid growth) and lag phases are traits associated with resilience after perturbations, however, these traits usually come at the cost of reduced resource use efficiency (23, 24). Similarly, also resistance traits can be associated with metabolic costs. Such costs may for instance be related to the strategy of generalists to process more environmental information than specialists and regulate their responses accordingly (25).

Microbial communities have been shown to allocate available resources either towards resistance or resilience, where oligotrophic environments favored resistant but slowly recovering communities while the opposite was true for nutrient-rich environments (e.g. Piton et al., 2020). However, in contrast to such resistance-resilience trade-off have oligotroph aquatic bacterial taxa like SAR11 been described as slow growing organisms with limited physiological and metabolic flexibility that are consequently sensitive to environmental change (27). Accordingly may nutrient limitation in some cases also select for taxa with simultaneous low resistance and low resilience. To address these conflicting observations, a recent study investigated trait-trait variations of the above highlighted genomic traits as markers for resilience and resistance from ~18,000 bacterial genomes to address trade-off between resistance and resilience traits (18). The observed trait-trait variations confirmed the hypothesized negative correlation (i.e. trade-off) of resistance versus resilience-related genomic traits only in prokaryotes with genomes >5 Mbp, which are typically found in soil environments. In contrast, for taxa with genome sizes that are typical for aquatic prokaryotes (i.e. <4 Mbp), resistance and resilience-related genomic traits correlated positively. This in silico analysis suggests that the addition of nutrients to aquatic environments would lead to a simultaneous increase of both, resistance and resilience.

Here, leveraging continuous cultivation experiments with complex aquatic bacterial communities, we address the hypothesis that exposure to multiple consecutive salinity pulse disturbances selects under sufficient nutrient supply for simultaneously resistant and resilient community members. We posit that this selection imposed by consecutive perturbations manifests in non-stochastic community re-assembly and reflects genomic traits associated with resistance and resilience. To unravel trade-offs between resistance and resilience and resource availability, we contrasted two levels of resource availability. Furthermore, we monitored bacterial biomass production and respiration to derive bacterial growth efficiency. Our findings highlight the importance of genomic traits in explaining community responses to consecutive perturbations. We provide experimental evidence for constraints of resource availability on resistance and resilience, which is critically important in light of multiple and simultaneous global changes.

## Results and Discussion

We performed a 41-days continuous culture (i.e., chemostat) experiment with repeated pulse disturbances and undisturbed control treatments crossed with two resource availability levels (Fig. 1). Weekly pulse disturbances were induced by adding a saturated NaCl solution, which resulted in a salinity increase of ~ 13 psu in the disturbed treatments (Fig. 1). The constant inflow of fresh medium washed out the added salt, thereby leading to salinity pulses. In total, the communities were exposed to 6 pulses. Resource availability was manipulated by two different sources of dissolved organic matter (DOM) representing low and high nutrient levels (L and HDOM). As expected, differences in nutrient availability were reflected in different cell abundances with 0.8 ± 0.4 × 10^6^ cells mL^-1^ (LDOM) and 2.1 ± 0.6 × 10^6^ cells mL^-1^ (HDOM), respectively. These values represent cell densities that are typical for oligotrophic to mesotrophic aquatic environments.

**Figure 1.**
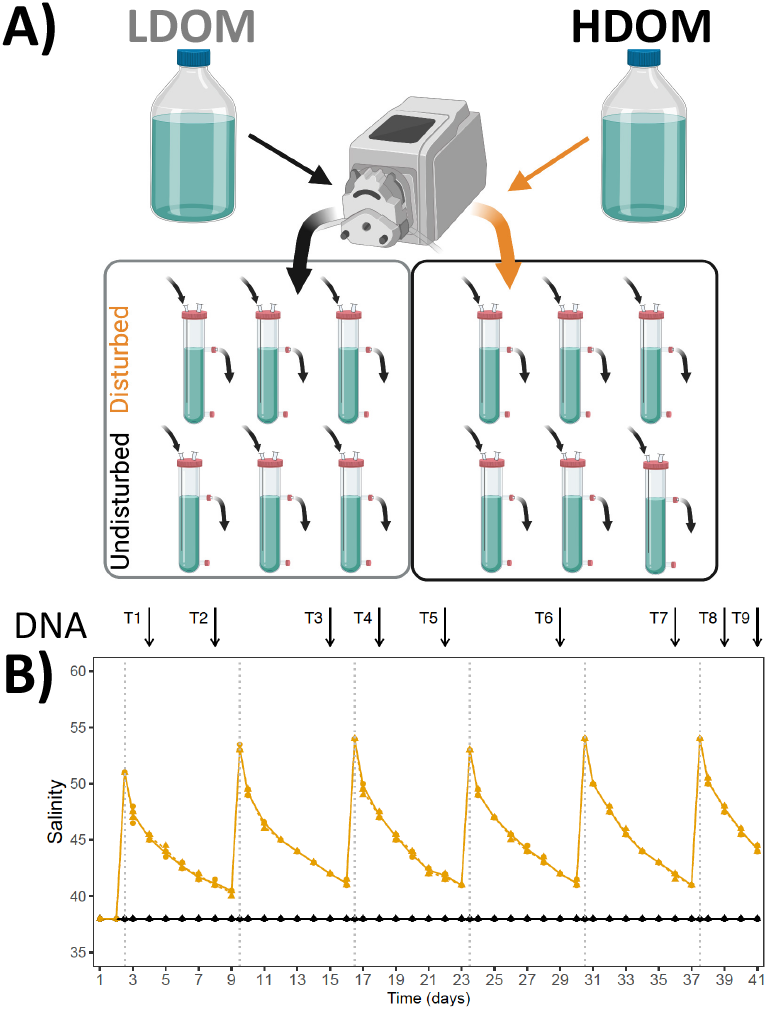
Experimental setup. **A)** Graphical schema summarizing the main experimental conditions, and **B)** salinity induced-pulse and the DNA sampling frequency.

We opted for a ‘super-diverse’ metacommunity as inoculum such that selection during experimental perturbations could act on a rich assemblage with complementary traits. The cultures were therefore inoculated with a metacommunity composed of microbial communities from several coastal Mediterranean marine and lagoon sites that differed concerning their disturbance history and eutrophication level (Fig. S1).

### Community composition and assembly

16S rRNA gene amplicon sequencing showed that early experimental stages were dominated by members of Flavobacteriales and Enterobacterales (Fig. S2). At intermediate stages, Rhodobacterales and Caulobacterales increased in abundance. At the end of the experiment, diverse taxonomies were observed even among replicates, including, for instance, members of the Caulobacterales, Sphingomonadales, Rhodospirillales, and Enterobacterales.

Multivariate Analysis of Variance (PERMANOVA) based on amplicon sequence variants (ASVs), suggested that time was the strongest structuring factor of community composition (F=51.9; P≤0.001). A pronounced taxon turnover was supported by Bray Curtis distance > 0.72 between the first and the last sample (T1 vs T9). Resource availability was the second strongest factor shaping bacterial communities (F=22.8, P≤0.001), followed by disturbance regime (F=2.5; P=0.03) (Fig. 2). Considerable divergence in community compositional patterns among replicated treatments, both at the ASV and order level (Fig. S2) indicated a pronounced stochastic compositional re-assembly component.

**Figure 2.**
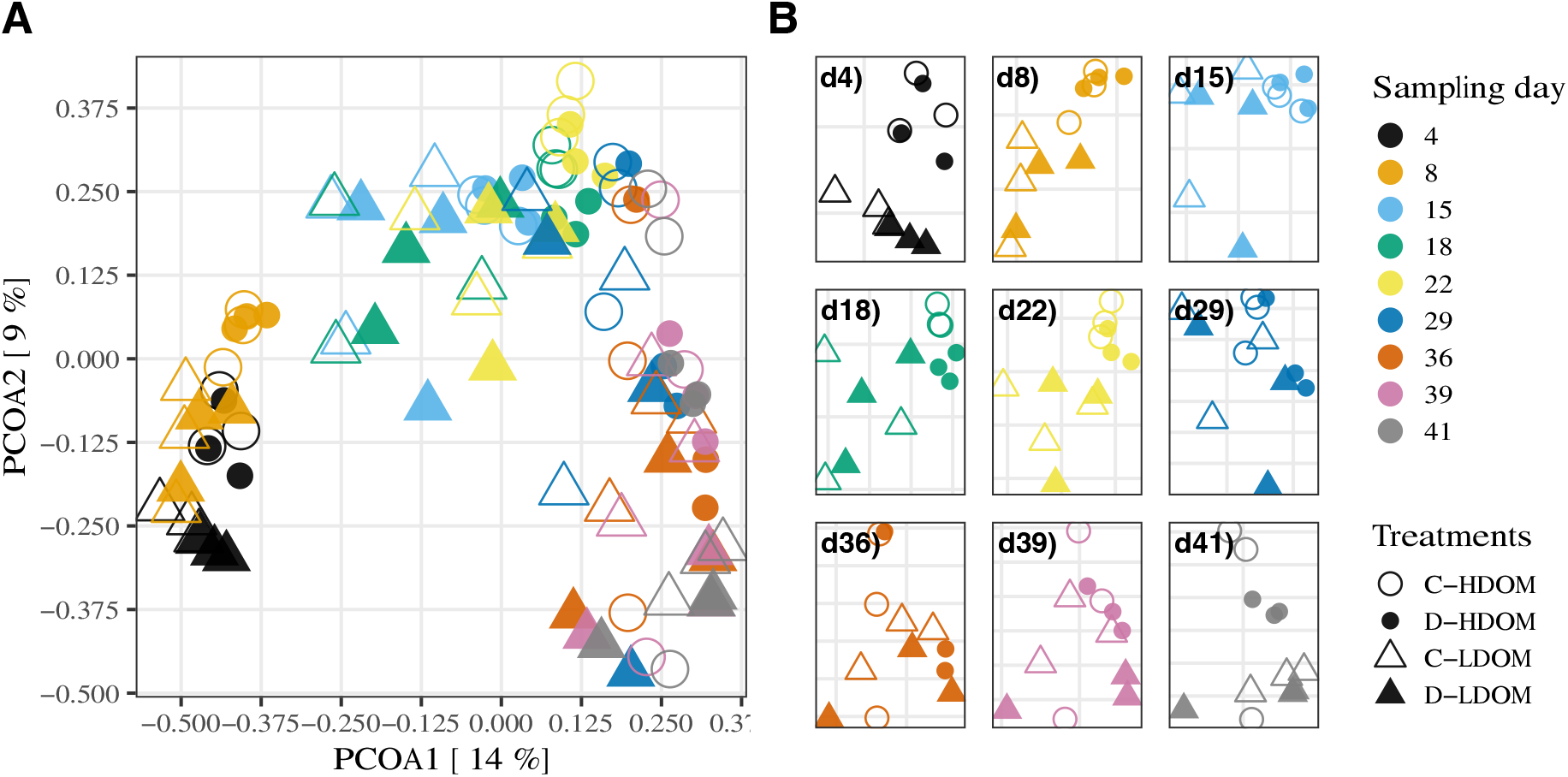
Community structures. A) Overview PCoA biplot including all data points (Bray-Curtis distances). B) PcoAs (Bray Curtis distances) for individual sampling days; axes of the individual plots are differently scaled as indicated by the grid lines.

Analyses of beta nearest taxon indices (bNTIs) pointed to a treatment-independent prevalence of stochastic rather than deterministic events during community re-assembly (Fig. S3). Our findings thereby contrast results from a continuous culture study in which periodical disturbance induced a shift towards a more deterministic assembly of microbial communities (28). However, another continuous culture experiment suggested that high species diversities in the starting communities increase the contribution of stochastic events during community assembly (29). Possibly, the highly diverse initial metacommunity inoculum contributed to stochastic re-assembly processes in our cultures.

Apart from stochasticity in community re-assembly, more deterministic selection may have shaped trait distribution. Both, bacterial life history traits as well as the above highlighted resistance and resilience-related genomic features have been shown to be phylogenetically conserved (18, 30). Phylogenetic distance-based community composition should accordingly reflect a greater impact of the disturbance regime compared to the purely compositional metric applied above. Indeed, PERMANOVA based on a phylogenetic instead of a compositional distance metric resulted in a comparably higher impact of the disturbance regime (F=5.0, P=0.006) (Fig. S4). This indicates that repeated pulse perturbations lead to a considerable trait-based similarity of communities with only moderate overlap in taxonomic composition.

### Genomic trait distributions and consequences on diversity patterns

To address the distribution of genomic traits, indicative of either resistance or resilience, trait values were extrapolated from databases of close relatives for individual ASVs. This allowed us to compute community-weighted means (CWMs, i.e. mean trait values weighted for ASV relative abundances) of genomic traits in individual communities. CWMs of all genomic traits with exception of genome size changed significantly over time and independently of the disturbance regime or resource availability (Fig. 3; Table 1). These temporal dynamics were particularly evident for resilience-related traits with a pronounced increase of minimal generation times that was extrapolated from codon usage bias (22). Likewise, a decrease of the RRN after the second experimental week. In line with these observations, high RRNs have been described during the early successional stages of microbial communities from different environments (31, 32).

**Figure 3.**
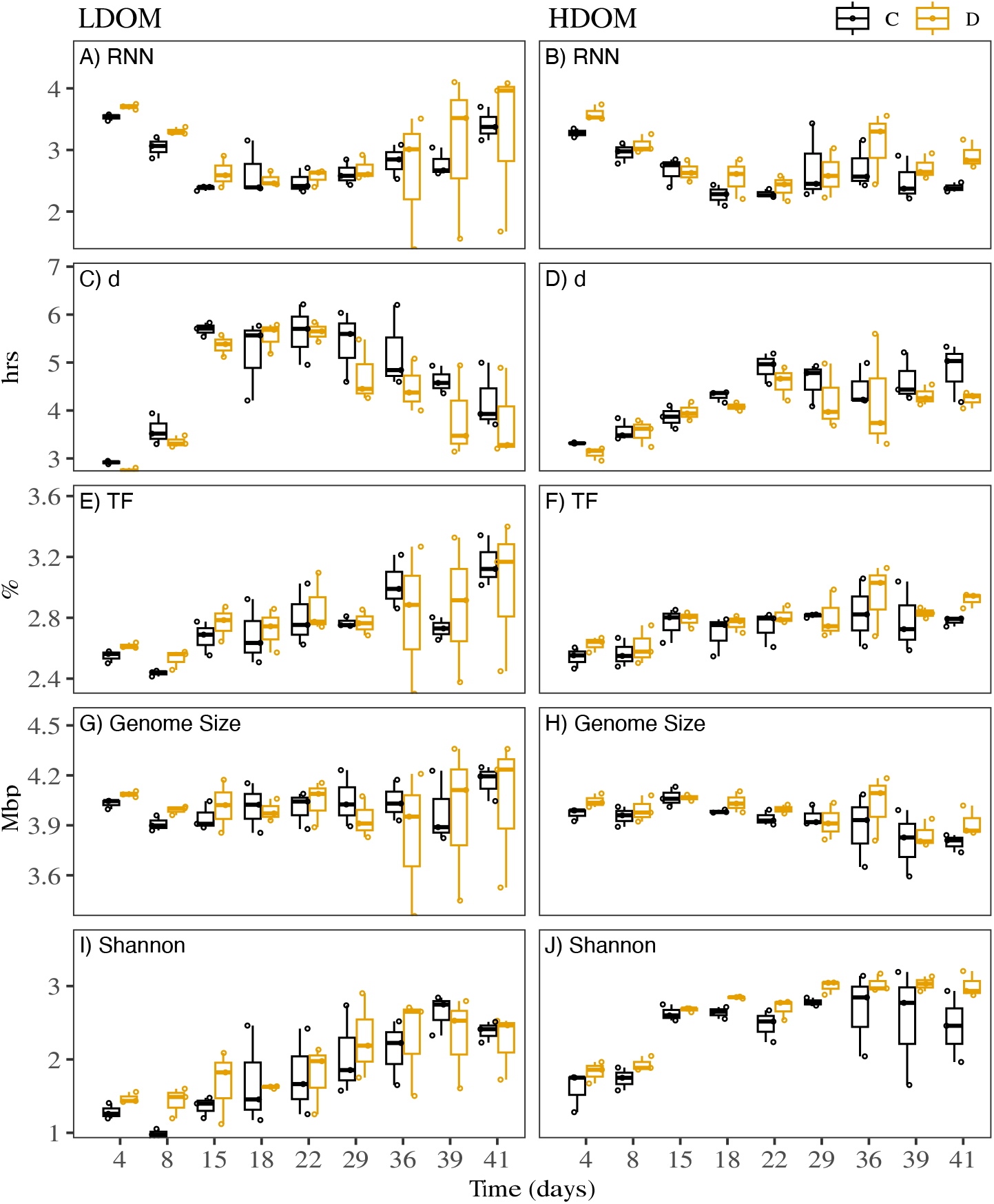
Boxplots displaying CWMs of genomic traits. LDOM and HDOM are displayed in the left and right panels, restively. **A, B)** RRN, **C, D)** d, **E, F)**, %TF **G, H)** genome size and **I, J)** Shannon diversity index.

**Table 1.**
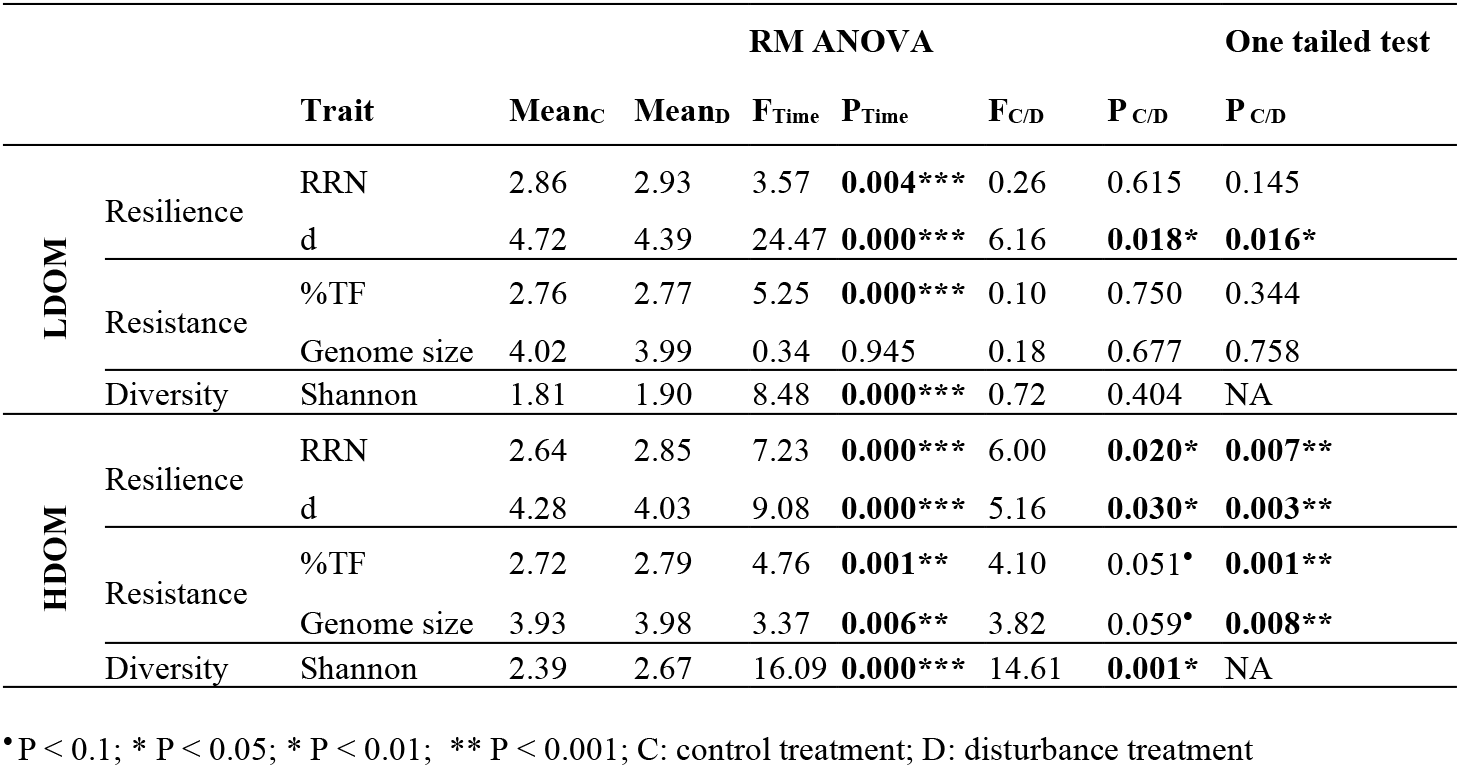
Results of statistical test to evaluate the effect of disturbance on genomic trait distributions.

Strikingly, besides the predominance of stochastic species-level assembly processes that occurred independent of the disturbance regime, consistent disturbance-driven effects were apparent for trait distributions. Disturbances impacted the distribution of genomic traits in dependence of resource availability, largely following our predictions: in the absence of resource limitation (HDOM) all evaluated genomic traits (genome size, %TF; RRN, minimal generation times) differed significantly between the disturbed and control treatments while suggesting a simultaneous increase of resistance and resilience in response to pulsed disturbances (Fig. 3, Table 1). In contrast, under limiting nutrient conditions (LDOM), only maximal generation time increased significantly in response to disturbances. However, also in the presence of resource constraints, no trade-offs between resistance and resilience were apparent (Fig. 3, Table 1). Taken together, our results indicate that generalist taxa with large genomes and elevated %TF only had competitive advantages under high resource availability, suggesting the existence of metabolic costs related to these traits.

Both, taxa resistant to environmental change and resilient taxa that rapidly recover after disturbances should benefit from a temporally variable environment. In line with this, the disturbance of communities with simultaneously high resistance and resilience has been predicted to lead to increased diversity (16). In agreement with these theoretical considerations, we found that in the absence of resource limitation, disturbances simultaneously increased resistance and resilience and resulted in significantly higher bacterial diversity compared to undisturbed controls. In contrast, in communities grown under resource limitation, disturbances did not result in increased resistance and species diversity did not differ from control treatments (Fig. 3I, Table 1).

### Community functional resistance

Both, a shift towards more resistant community members, but also increased diversity have been debated as mechanisms that can increase community-level functional resistance to disturbances (10, 17). In order to evaluate the functional consequences of species and trait selection processes, we measured bulk community functional resistances to the weekly induced pulse disturbances. A resistance index was computed by quantifying the relative change of heterotrophic bacterial production (BP), respiration, and growth efficiency (BGE) before and after each disturbance compared to the corresponding activity change in the respective control treatments. We found that without resource limitation (HDOM), bacterial communities exhibited significantly higher resistance of BP than in communities exposed to resource limitation (LDOM)(Fig. 4A, Table 2). Noticeably, patterns of bacterial biomass production were manifested in cell abundance throughout the experiment. Specifically, in communities grown under resource limitations (LDOM) salinity disturbances lead to low resistance of biomass production causing a significant decrease in cell abundance. In contrast, under alleviated resource conditions (HDOM), biomass production was highly resistant and cell numbers remained unaffected by disturbances (Table 2, Fig. 4D). These findings highlight the role of resource availability for the resistance of aquatic microbial communities. We posit that resource abundance can alleviate metabolic costs associated with resistance (adaptability) and fundamentally alter functional consequences of perturbations. This has important consequences considering multiple and simultaneous perturbations associated with global change, including wide-spread eutrophication.

**Figure 4.**
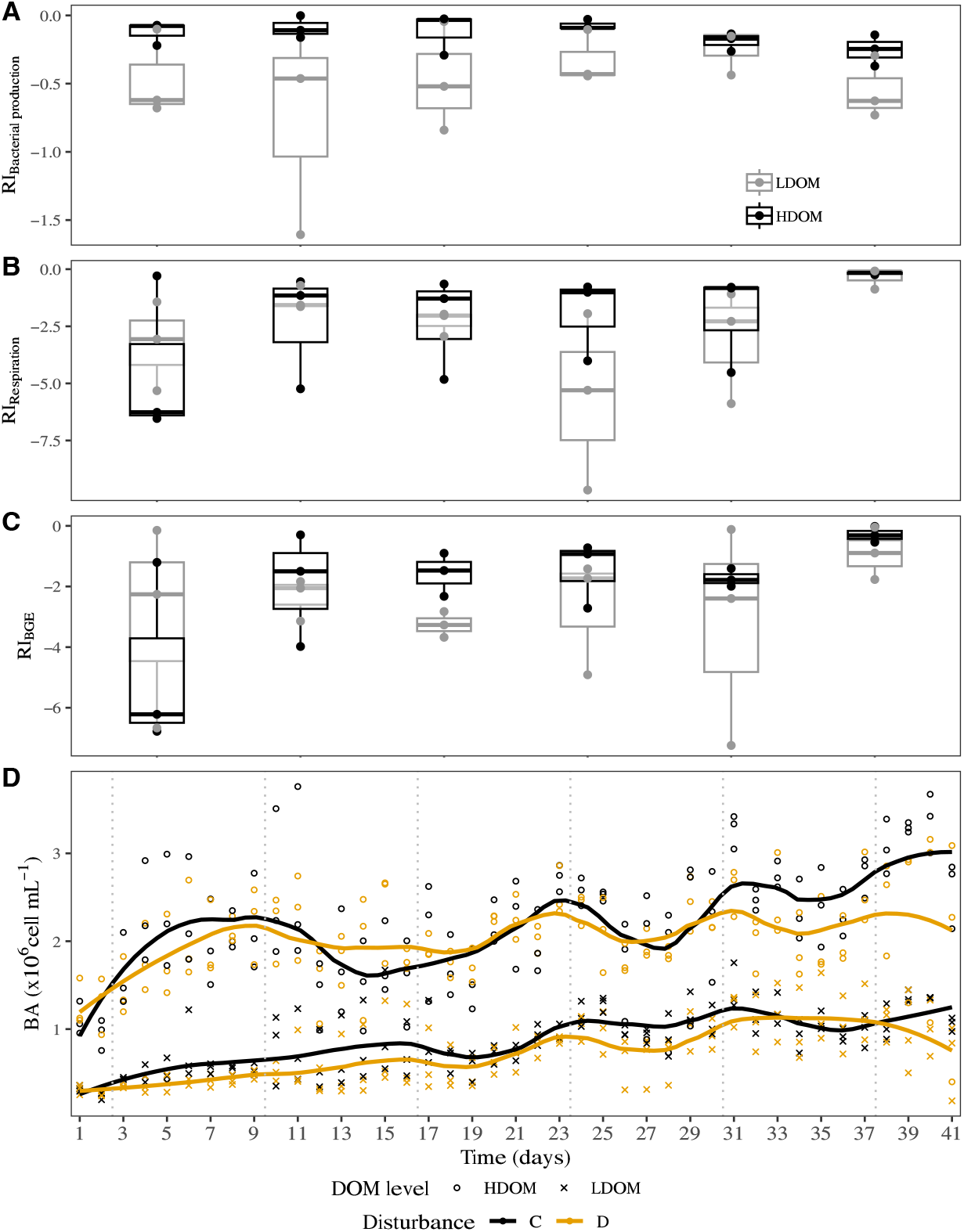
Community functional parameters. Resistance indices (gray: LDOM, black: HDOM) for **A)** BP, B) respiration and C) BGE. **D)** Bacterial abundance, lines indicate locally fitted values (loess smoothing) under each disturbance and DOM regime.

**Table 2.** Results of statistical tests to evaluate the effect of nutrient availability to functional resistance

| RIs Bulk<br>functioning | RM ANOVA |  |  |  | MLM<sub>LDOM</sub> | MLM<sub>HDOM</sub> |
| --- | --- | --- | --- | --- | --- | --- |
|  | F<sub>Time</sub> | P<sub>Time</sub> | F<sub>DOM</sub> | P<sub>DOM</sub> | Slope | Slope |
| BP | 0.56 | 0.726 | 10.83 | **0.003\*** | 0.04 | -0.03 |
| Respiration | 2.13 | 0.098* | 0.41 | 0.527 | 0.17 | **0.63\*** |
| BGE | 1.72 | 0.170 | 0.92 | 0.347 | 0.24 | **0.66\*** |

\* P < 0.05

Interestingly, we could not detect significant temporal trends in BP resistance (Fig. 4A, Table 2). Apparently, consecutive perturbations did not select for increasingly more functionally resistant community members. Instead, the observed differences between the HDOM and LDOM treatments seemed inherent to the respective communities grown under different nutrient regimes. Differences in cell densities between disturbed versus control LDOM regimes already after the first disturbance (Fig. 4D) as well as non-consistent temporal patterns in resistance trait distributions (Fig. 3G, H) reflect a roughly constant BP resistance over time. This is particularly noteworthy in consideration of the pronounced successional turnover, indicating that resource limitation may exert stronger controls on the functional consequences of perturbations than community composition or diversity. This is important concerning potential management strategies.

Temporal patterns of resistance for respiration, as well as BGE, clearly differed from those observed for BP, by suggesting at least in the HDOM incubations a resistance increase over time in response to consecutive disturbances. Furthermore, in contrast, to bacterial biomass production, neither respiration nor growth efficiency differed significantly between treatments with (LDOM) and without (HDOM) resource limitation (Table 2). We attribute this to the contribution of maintenance metabolism to bacterial respiration, which is independent of resource availability.

Taken together, our analyses point to different mechanisms that determine resistance of biomass production and respiration. The responses of resistance and resilience traits, diversity, biomass production and growth point towards a pivotal role of resource availability in determining the outcomes of perturbations. We suggest that in the presence of resources, modulations of growth can counteract the effects of perturbations, leading to apparent resistance and resilience despite large taxa turnover. On the other hand, when resources are limiting, minimum requirements for cellular respiration (reflected in BGE), putatively related to protein and RNA repair, seem to constrain the response space for bacterial communities facing perturbations.

### Conclusions

This work provides empirical evidence for the role of resource availability in determining the compositional and functional responses of bacterial communities to perturbations. Our results indicate community re-assembly towards resistant and resilient taxa with consequences for community diversity and functioning. However, this capacity to re-assemble was restricted by nutrient availability. While stochasticity during community re-assembly was important, we found consistent patterns of genomic trait distributions, diversity patterns and functional characteristics. Our findings, therefore, suggest that selection occurred on traits rather than on individual populations, thereby leading to alternative compositional solutions with similar functionality. This sheds new light on the notion of prevalent functional redundancy in microbial communities. Better understanding the functional consequences and underlying mechanisms of microbial community resistance and resilience will be important in light of increasing frequencies, concomitance and magnitude of multiple global change perturbations to ecosystems, including coastal marine areas.

## Material and Methods

### Starting communities and culture media

Microbial inocula for the continuous culture experiment were sampled from several sites in the south of the Gulf of Lyon, South France that feature contrasting environmental variability (Fig. S1; Table S1). Larger organisms, such as protists were excluded by a pre-filtration step. Communities for these sites were conserved by applying a protocol for cryopreservation (33) and cryo-aliquots from all sites were pooled in culture media at the experiment start. The cultures media were based on artificial seawater (ASW; salinity: 38 psu, pH: 8; Eguchi *et al*., 1996) and amended with DOM supplements as sole carbon, nitrogen, and phosphorus source, while no vitamins were added. Trace metals, Fe, and EDTA were added 100-fold less concentrated than originally published.

Two complex DOM supplements that supported the growth of cell densities as typically found in oligo-to mesotrophic conditions (LDOM) or eutrophic conditions (HDOM) were prepared from different aquatic environments (Table S2) as described elsewhere (33). HDOM media were additionally amended with yeast extract (0.28 mg L^-1^; Sigma–Aldrich, St. Louis, MO, United States).

### Experimental design

Overall, the setup of the continuous culture system resembled that of an earlier continuous culture study (Baho et al., 2012). We combined triplicate treatments of two disturbance regimes (undisturbed control, weekly disturbance +13 psu) with two different resource availability treatments (LDOM, HDOM). We started the long-term experiment with a preculture that was set up in batch modus before turning to the continuous mode and using the inoculum detailed above. The applied flow rate of 70 μL min^-1^ was set to approximate generation time measured for prokaryote communities from the Mediterranean sea (~2.75 days, 36). The incubations were kept at 18°C in dark during the entire experiment. In total, 6 pulse disturbances were applied within 41 days of continuous flow mode (Fig. 1; Table S3).

We tested the medium via flow cytometry (FC) regularly for potential contamination after it had passed the tubing system just before the inlet into the continuous culture. While occasional contaminations were detected, these never exceeded 10^5^ cells per mL^-1^ and were minor compared to cell concentrations in the vessels (<6.6%, Table S4). These sporadically occurring contaminations are therefore unlikely to explain the strikingly dissimilar communities among replicates from all treatments after the second experimental week (Fig. S2).

### Community assembly

Community aliquots for DNA extraction and downstream metabarcoding were sampled at least weekly from the continuous culture outflow. In total, we obtained DNA samples from 9 sample days (Fig. 1, Table S3). Filters for DNA extractions were stored at −30°C until further processing.

DNA extractions were performed using a QIAmp DNA Mini Kit (QIAGEN, Hilden, Germany) and sent for 16s rRNA gene amplicon sequencing using the primers pair 515yf-926r (37). Sequence processing was performed using the DADA2 for R (38). A total of 1447 ASVs were identified and were taxonomically assigned using the Genome Taxonomy Database (GTDB) (39). The taxonomic and phylogenetic compositional structure of the communities were assessed using Bray-Curtis distances or the pairwise abundance weighted UniFrac metric (Lozupone and Knight, 2005), respectively. To evaluate the impact of nutrient and disturbance regimes as well as time on community assembly we performed a PERMANOVA.

### Genomic trait distributions and species diversity

Genomic traits (resistance-related traits: genome size, %TF; resilience-related traits: RRN, generation time delineated from codon usage biases) were predicted for each ASV using the hidden state prediction option included in the PICRUSt2 software v2.4.2 (41) and using the trait values published elsewhere (18, 42). CWMs of genomic trait data in each sample were computed by multiplying the predicted trait values for each ASV by its relative abundance and adding up these weighted values in each sample. The Shannon diversity index was computed from the ASV compositional data to estimate species diversity. We performed two-way repeated-measurements analyses of variances (rmANOVAs) to assess the effect of time and the disturbance regime on bacterial abundances, Shannon diversity, and genomic traits in each of the DOM regimes. In the case of genomic traits, we also performed a one tailed paired t-test on the mean values of the genomic traits to test the specific a-priory hypothesis that resistance and resilience values would both increase (and not decrease) in response to pulsed salinity disturbances.

### Community functional measurements

Cell growth in the continuous culture was estimated by measuring cell densities via FC as detailed elsewhere (43).

Functional resistance of the continuous culture was estimated weekly by measuring the below-described bulk community functional rates before and 1 hour after each induced disturbance in the disturbance treatments and simultaneously also in the controls (Table S3). BP was assessed via ^3^H-leucine incorporation (44). Respiration was quantified as the oxygen consumption in 5 mL glass using a SensorDish reader (PreSens, Regensburg, Germany). BGE was estimated by dividing BP by the sum of BP and respiration.

We applied log-response ratios (lnR, Hillebrand and Gurevitch, 2016) to quantify the relative change of the functional rate F before and after the pulse disturbance in the disturbed communities (lnR_D_) and in the corresponding controls where no disturbance was introduced (lnR_C_). We considered the difference between both ratios as the resistance index for function F (RI_F_) which is similar to the effect size measurement published previously by Osenberg et al (1997).

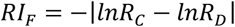

The larger the deviation of RI_F_ from zero, the lower/higher the functional resistance/sensitivity.

To test the impact of culture conditions over time on bulk community functional resistances we performed two-way rmANOVAs on the resistance indices (RI_F_) considering time and DOM level. We additionally fitted a linear mixed model (LMM) on the RI by DOM level to evaluate the direction of potentially detected trends over time for each of both DOM conditions separately.

## Data Availability

R-scripts and unix shell scripts used for the data analyses have been published on GitHub (https://github.com/angelrainf/chemo.disturbances). The sequence data for this study have been deposited in the European Nucleotide Archive (ENA) at EMBL-EBI under accession number PRJEB55672.

## ACKNOWLEDGMENTS

This study was financed by the German Science Foundation (DFG) granted to SB (BE 5937/2-1). ARF was supported by a scholarship from the Chilean National Agency for Research and Development (ANID) / Scholarship Program / DOCTORADO BECAS CHILE/2017 – 72180448. GPM was supported by the doctorate scholarship program of the Coordination for the Improvement of Higher Education Personnel (CAPES), Brazil, and the above-mentioned DFG funding. The bioinformatics work was supported by the BMBF-funded de.NBI Cloud within the German Network for Bioinformatics Infrastructure (de.NBI) (031A532B, 031A533A, 031A533B, 031A534A, 031A535A, 031A537A, 031A537B, 031A537C, 031A537D, 031A538A). We thank the Service d’Observation en Milieu Littoral (SOMLIT-Banyuls-sur-Mer) for providing data, and the facilities to collect water samples. We thank the BIO2MAR platform for the access and technical support in their molecular biology facilities. We thank Olivier Crispy, Jenny Jeschek, and Madleen Dierken for their help in the analysis of dissolved organic compounds and nutrients. We also thank Fernanda González for her support during sampling activities.

## Notes

https://github.com/angelrainf/chemo.disturbances

https://www.ebi.ac.uk/ena/browser/view/PRJEB55672

### Competing Interest Statement

The authors have declared no competing interest.

## REFERENCES

1. J. F. Johnstone, et al., Changing disturbance regimes, ecological memory, and forest resilience. Frontiers in Ecology and the Environment 14, 369–378 (2016).

2. D. A. Smale, et al., Marine heatwaves threaten global biodiversity and the provision of ecosystem services. Nat. Clim. Chang. 9, 306–312 (2019).

3. D. Gampe, et al., Increasing impact of warm droughts on northern ecosystem productivity over recent decades. Nat. Clim. Chang. 11, 772–779 (2021).

4. L. Samaniego, et al., Anthropogenic warming exacerbates European soil moisture droughts. Nature Clim Change 8, 421–426 (2018).

5. M. C. Kirchmeier-Young, X. Zhang, Human influence has intensified extreme precipitation in North America. Proceedings of the National Academy of Sciences 117, 13308–13313 (2020).

6. B. Sun, et al., Experimental warming reduces ecosystem resistance and resilience to severe flooding in a wetland. Science Advances 8, eabl9526 (2022).

7. D. Breitburg, et al., Declining oxygen in the global ocean and coastal waters. Science 359, eaam7240 (2018).

8. S. S. Kaushal, et al., Freshwater salinization syndrome on a continental scale. Proceedings of the National Academy of Sciences 115, E574–E583 (2018).

9. M. Le Moal, et al., Eutrophication: A new wine in an old bottle? Science of The Total Environment 651, 1–11 (2019).

10. F. Isbell, et al., Biodiversity increases the resistance of ecosystem productivity to climate extremes. Nature 526, 574–577 (2015).

11. C. R. Biggs, et al., Does functional redundancy affect ecological stability and resilience? A review and meta-analysis. Ecosphere 11, e03184 (2020).

12. R. Cavicchioli, et al., Scientists’ warning to humanity: microorganisms and climate change. Nat Rev Microbiol 17, 569–586 (2019).

13. A. Shade, et al., Fundamentals of microbial community resistance and resilience. Front. Microbiol. 3, 417 (2012).

14. B. S. Griffiths, L. Philippot, Insights into the resistance and resilience of the soil microbial community. FEMS Microbiology Reviews 37, 112–129 (2013).

15. A. Rain-Franco, N. Mouquet, C. Gougat-Barbera, T. Bouvier, S. Beier, Niche breadth affects bacterial transcription patterns along a salinity gradient. Molecular Ecology 31, 1216–1233 (2022).

16. D. G. Nimmo, R. M. Nally, S. C. Cunningham, A. Haslem, A. F. Bennett, Vive la resistance: reviving résistance for 21st century conservation. Trends in Ecology & Evolution 30, 516–523 (2015).

17. M. G. Matias, M. Combe, C. Barbera, N. Mouquet, Ecological strategies shape the insurance potential of biodiversity. Front. Microbiol. 3, 432 (2013).

18. S. Beier, J. Werner, T. Bouvier, N. Mouquet, C. Violle, Trait-trait relationships and tradeoffs vary with genome size in prokaryotes. Frontiers in Microbiology 13(2022).

19. I. Kostadinov, et al., Quantifying the effect of environment stability on the transcription factor repertoire of marine microbes. Microbial Informatics and Experimentation 1, 9 (2011).

20. P. Bentkowski, C. Van Oosterhout, T. Mock, A Model of Genome Size Evolution for Prokaryotes in Stable and Fluctuating Environments. Genome Biol. Evol. 7, 2344–2351 (2015).

21. J. A. Klappenbach, J. M. Dunbar, T. M. Schmidt, RRNA operon copy number reflects ecological strategies of bacteria. Appl. Environ. Microbiol. 66, 1328–1333 (2000).

22. J. L. Weissman, S. Hou, J. A. Fuhrman, Estimating maximal microbial growth rates from cultures, metagenomes, and single cells via codon usage patterns. PNAS 118(2021).

23. N. Fierer, M. A. Bradford, R. B. Jackson, Toward an ecological classification of soil bacteria. Ecology 88, 1354–1364 (2007).

24. B. R. K. Roller, S. F. Stoddard, T. M. Schmidt, Exploiting rRNA operon copy number to investigate bacterial reproductive strategies. Nat Microbiol 1, 1–7 (2016).

25. S. R. X. Dall, I. C. Cuthill, The information costs of generalism. Oikos 80, 197–202 (1997).

26. G. Piton, et al., Using proxies of microbial community-weighted means traits to explain the cascading effect of management intensity, soil and plant traits on ecosystem resilience in mountain grasslands. Journal of Ecology 108, 876–893 (2020).

27. D. L. Kirchman, “Growth Rates of Microbes in the Oceans” in Annual Review of Marine Science, Vol 8, C. A. Carlson, S. J. Giovannoni, Eds. (Annual Reviews, 2016), pp. 285–+.

28. M. S. Gundersen, I. A. Morelan, T. Andersen, I. Bakke, O. Vadstein, The effect of periodic disturbances and carrying capacity on the significance of selection and drift in complex bacterial communities. ISME COMMUN. 1, 1–9 (2021).

29. J. M. Ayarza, L. Erijman, Balance of Neutral and Deterministic Components in the Dynamics of Activated Sludge Floc Assembly. Microb. Ecol. 61, 486–495 (2011).

30. S. P. Blomberg, T. Garland, A. R. Ives, Testing for phylogenetic signal in comparative data: Behavioral traits are more labile. Evolution 57, 717–745 (2003).

31. R. Niederdorfer, K. Besemer, T. J. Battin, H. Peter, Ecological strategies and metabolic trade-offs of complex environmental biofilms. npj Biofilms Microbiomes 3, 1–6 (2017).

32. J. Guittar, A. Shade, E. Litchman, Trait-based community assembly and succession of the infant gut microbiome. Nat Commun 10, 512 (2019).

33. A. Rain-Franco, G. P. de Moraes, S. Beier, Cryopreservation and Resuscitation of Natural Aquatic Prokaryotic Communities. Frontiers in Microbiology 11, 3633 (2021).

34. M. Eguchi, et al., Responses to Stress and Nutrient Availability by the Marine Ultramicrobacterium Sphingomonas sp. Strain RB2256. Appl. Environ. Microbiol. 62, 1287 (1996).

35. D. L. Baho, H. Peter, L. J. Tranvik, Resistance and resilience of microbial communities - temporal and spatial insurance against perturbations. Environ. Microbiol. 14, 2283–2292 (2012).

36. M. Landa, et al., Phylogenetic and structural response of heterotrophic bacteria to dissolved organic matter of different chemical composition in a continuous culture study. Environ Microbiol 16, 1668–1681 (2014).

37. A. E. Parada, D. M. Needham, J. A. Fuhrman, Every base matters: assessing small subunit rRNA primers for marine microbiomes with mock communities, time series and global field samples. 18, 1403–1414 (2016).

38. B. J. Callahan, et al., DADA2: High-resolution sample inference from Illumina amplicon data. 13(2016).

39. D. H. Parks, et al., A standardized bacterial taxonomy based on genome phylogeny substantially revises the tree of life. Nature Biotechnology 36, 996–1004 (2018).

40. C. Lozupone, J. R. Knight, UniFrac: a new phylogenetic method for comparing microbial communities. Applied and Environmental Microbiology 71, 8228–8235 (2005).

41. G. M. Douglas, et al., PICRUSt2 for prediction of metagenome functions. Nature Biotechnology 38, 685–688 (2020).

42. S. F. Stoddard, B. J. Smith, R. Hein, B. R. K. Roller, T. M. Schmidt, rrnDB: improved tools for interpreting rRNA gene abundance in bacteria and archaea and a new foundation for future development. Nucleic Acids Res. 43, D593–D598 (2015).

43. D. Marie, N. Simon, L. Guillou, F. Partensky, D. Vaulot, “Flow Cytometry Analysis of Marine Picoplankton” in In Living Color: Protocols in Flow Cytometry and Cell Sorting, Springer Lab Manuals., R. A. Diamond, S. Demaggio, Eds. (Springer, 2000), pp. 421–454.

44. D. Smith, F. Azam, A simple economical method for measuring bacterial protein synthesis rates in seawater using 3H-Leucine. Marine Microbial Food Webs 6, 107–114 (1992).

45. H. Hillebrand, J. Gurevitch, “Meta-Analysis and Systematic Reviews in Ecology” in ELS, John Wiley & Sons Ltd, Ed. (John Wiley & Sons, Ltd, 2016), pp. 1–11.

46. C. W. Osenberg, O. Sarnelle, S. D. Cooper, Effect Size in Ecological Experiments: The Application of Biological Models in Meta-Analysis. The American Naturalist 150, 798–812 (1997).

