## Supplementary text and figures for "The Cost of Adaptability: Resource Availability Constrains Functional Stability Under Pulsed Disturbances"

**This PDF file includes:**

Supporting text
Figures S1 to S4
SI References

### Supporting Information Text

#### Material and Methods

#### Starting communities

Microbial inocula for the continuous culture experiment were obtained from cryopreserved microbial community aliquots that had been prepared as detailed earlier (Rain-Franco et al., 2021). In short, microbial communities were prefiltered using a 0.8 µm pore size membrane filter to exclude larger organisms, such as protists, and subsequently concentrated by filtering om a 0.2 µm pore size membrane filter. The concentrated cells on the filter were merged in 1 ml culture medium containing 5% DMSO as cryo-protectant and after 15 minutes at 4°C shock frozen in liquid nitrogen. These community aliquots were conserved at -80°C until the experiment start.

The aquatic environments from which the community aliquots had been sampled comprised several sites in the south of the Gulf of Lyon, South France. The sites including the Mediterranean field station SOLA and the coastal lagoons La Palme and Gruissan are characterized by contrasting environmental variability (Fig. S1; Table S1). Cryopreserved community aliquots from these sites were pooled to obtain a ‘super-diverse’ metacommunity with the aim that selection mechanisms during downstream experimental treatments could act on a possibly large collection of species with complementary traits.

#### Culturing media

Cultures media in this study were based on artificial seawater (ASW; Eguchi *et al.*, 1996) with a salinity of 38 g $L^{-1}$ and pH 8. Trace metals, Fe, and EDTA were added 100-fold less concentrated than originally published. The ASW was amended with DOM supplements as sole carbon, nitrogen, phosphorus source, and no vitamins were added.

DOM supplements were prepared from particular material retained after filtration of water from 7 different aquatic environments that differed in their trophic status (Table S2) as described elsewhere (Rain-Franco et al., 2021). Based on several pre-tests, we combined these DOM supplements into low and high nutrient DOM supplements (referred to as LDOM and HDOM, respectively, Table 2) that supported the growth of cell densities as typically found in oligo- to mesotrophic conditions (LDOM) or eutrophic conditions (HDOM). HDOM media were additionally amended with yeast extract (0.28 mg $L^{-1}$; Sigma–Aldrich, St. Louis, MO, United States) to add a highly labile DOM compound. Complex and diverse DOM sources as the DOM supplements for LDOM and HDOM media have been discussed to support the assembly of diverse prokaryotic communities. The preparation of concentrated LDOM and HDOM supplements allowed us further to repeatedly and reproducibly prepare large volumes of media that were necessary for the long-term continuous culture experiment.

### Experimental design

We started the long-term experiment with a preculture that was set up in batch modus. For this purpose cryopreserved communities from SOLA, Gruissan, and La Palme were resuscitated in 6 L ASW to which all 7 DOM supplements were added, each of them in half concentration compared to LDOM and HDOM media used later. This preculture was incubated for 3 days at 18°C under agitation by a magnetic stirrer. Subsequently, this preculture was split into two volumes. We added another half portion of either the LDOM or the HDOM media supplements to each of the volumes, respectively, except for the yeast extract, and maintained these cultures for 3 more days in batch condition before distributing them into the continuous system. After 6 days in batch mode, 300 ml of both precultures were distributed into 6 reactor vessels, respectively (Fig. 1). The inflow of LDOM and HDOM media was activated to fill the vessels with a volume of 400 ml and to start the continuous mode. From the 6 chemostats assigned to each DOM condition, 3 were used as control and 3 for the pulse disturbance treatment. The medium was pumped from 6 × 2 L bottles (3 containing LDOM and 3 containing HDOM medium) into the culture vessels using a peristaltic pump (ISMATEC, Cole-Parmer GmbH, Wertheim, Germany), while each bottle fed one control and one disturbance, respectively. The applied flow rate of 70 $\mu L\ min^{-1}$ was set to approximate generation time measured for prokaryote communities from the Mediterranean sea (~2.75, Landa et al., 2014). The cultures were mixed constantly with magnetic stirrers and ventilated with venting filters (0.2 µm, 64 mm diameter, Midisart® 2000, Sartorius, Göttingen, Germany) attached to the lids. Syringes (50 ml) were connected to the culture vessels via stainless steel needles and were used either to add saturated salt solutions to the disturbance treatments or to sample small volumes for flow cytometry (FC), salinity, and functional rate measurements. Water volumes of approximately 200 mL DNA extractions were sampled for downstream from the continuous culture's outflow from 2 consecutive days. Overall, the setup of the continuous culture system resembled that used in an earlier continuous culture study (Baho et al., 2012). We further added 2 polyethylene biochips (Mutag BioChip 25™, MUTAG, Chemnitz, Germany) into each vessel to provide a surface for biofilm formation. Cells attached to the biofilm that are not washed out can recolonize the medium and the biofilm on the Mutag BioChip 25™ may therefore serve as a spatial refuge. We decided to add these chips to lower the risk of a potential collapse of the continuous cultures during the long-term experiment.

After filling the reactor vessels, the cultures were stabilized in continuous mode for 2 days before the first pulse disturbance. Salt pulse disturbances were applied once per week to the disturbance treatments by adding ~18 ml of a saturated NaCl solution after the same volume was removed from the reaction vessels (+13 $gl^{-1}$ NaCl). The exact added volume was calculated each time and for each disturbance treatment vessel from the salinity measured in the respective vessel just before the addition of the salinity pulse to avoid a potential continuous raise of the salinity baseline and ensure that the final salinity after each peak did not exceed a salinity of ~54 $g\ L^{-1}$ NaCl. As a consequence of the continuous flow mode, the salt was diluted and after 7 days before the next pulse disturbance was introduced, the baseline salinity was approximately reached (Fig. 1A). In total, 6 pulse disturbances were applied within 41 days of continuous flow mode (Fig. 1A; Table S3).

The incubations were kept at 18°C in dark during the entire experiment. All components of the continuous culture system were autoclaved before the experiment start. We exchanged the medium and the tubing system every 3-4 days to avoid potential contamination. We further tested the medium regularly for potential contamination after it had passed the tubing system just before the inlet into the continuous culture, via FC. While during some occasions contaminations were detected, these never exceeded $10^5$ cells per $mL^{-1}$ (Table S4). In case of an indicated contamination, the whole tubing system and medium for the affected culture vessels were replaced with newly sterilized material.

### Community assembly

Samples for DNA extraction and downstream metabarcoding were taken from the continuous culture outflow and filtered onto 0.22 µm filters (cellulose filters, Millipore, MA, United States) (Table S3). To obtain a sufficiently large water volume, a sterile water bottle was attached to the outflow approximately 48 h before sampling. Sampling was performed at least once per week. In week 3, an additional set of samples was obtained as the continuous flow had accidentally stopped for some hours in two vessels (vessel 2 and 5; Fig. S3). Also, in week 6 two additional sets of DNA were sampled (Table S3). In total, we obtained DNA samples from 9 sample days (Fig. 1B). Filters for DNA extractions were stored at -30°C until further processing.

DNA extractions were performed using a QIAmp DNA Mini Kit (QIAGEN, Hilden, Germany) with an initial bead-beating step in ATL buffer using a FastPrep-24™ 5G (MP Biomedical, California, USA). The concentration and quality of the eluted DNA were tested using a DS11-FX+ microvolume spectrophotometer (DeNovix, Delaware, USA).
DNA samples from the continuous culture were sent for 16s rRNA gene amplicon sequencing (300 base pairs paired-end read, Illumina Miseq V3) using the primers pair 515yf-926r (Parada et al., 2016). Libraries were demultiplexed by the sequencing company (LGC Genomics GmbH) using the BCL2FASTQ software v2.17.1.14 that removed reads with length <100 bases and performed primer clipping (up to 3 mismatches per primer). Sequences were further processed using the DADA2 software for R (Callahan et al., 2016) by slightly modifying the standard pipeline (truncLen=c(250,180); Rain-Franco et al., 2021). A total of 1678 amplicon sequence variants (ASVs) were identified across samples. Retrieved ASVs were taxonomically assigned using the Genome Taxonomy Database (GTDB) (Parks et al., 2018). Read count data were rarefied to the minimum number of reads obtained across the samples (9,412 reads, Table S5 for all downstream data depending on community compositional data (beta diversity and alpha diversity dependent analyses, genomic trait distributions).

### Beta nearest taxon index (bNTI)

We computed the bNTIs to evaluate assembly mechanisms (Webb, 2000; Stegen et al., 2012) in response to the disturbance regimes under the different DOM levels. The bNTI evaluates whether the phylogenetic similarity between a pair of two samples was significantly lower or higher than expected by chance relative to a reference species pool. A phylogenetic similarity surpassing the theoretical expectation (bNTI >2) indicates the prevalence of variable deterministic assembly processes during community succession. A phylogenetic similarity below the theoretical expectation (bNTI< -2) indicates the prevalence of homogeneous deterministic assembly processes during community succession. Finally, a bNTIs between -2 and 2 indicates that community assembly is driven by stochastic processes rather than deterministic processes. NTIs were calculated via the iCAMP R-package (v1.3.4, Ning et al., 2020). All NTIs were estimated relative to the species pool in the incubations under the same DOM regime sampled during a single time point (=6 samples). This reference species pool was chosen because we aimed to focus on assembly mechanisms due to the disturbance regime for each of the DOM levels separately while excluding successional aspects.

### Genomic trait distributions and species diversity

Genomic traits that are indicative of the life history of prokaryotes feature significant phylogenetic signals and can be extrapolated from closely related relatives via taxonomic marker genes of uncultured species in a community (Beier et al., 2021). Genomic traits were predicted for each ASV using the hidden state prediction option included in the PICRUSt2 v.2.4.2, (Douglas et al., 2020) . This was done based on recently published trait values of strains included into the default PICRUSt2 reference tree for generation time, percent transcription factors (%TF) and genome size (Beier et al., 2021). However, due to possibly problematic RRN values of the internal PICRUSt2 RRN reference database (Beier et al., 2021) we instead predicted RRN using trait values available via the Ribosomal RNA Operon Copy Number Database (rrnDB, Stoddard et al., 2015). For this purpose, rrnDB 16s rRNA gene sequences (rrnDB v5.7) were aligned with the MUSCLE software (Edgar, 2004) and used to construct a phylogenetic tree via the FastTree software (vs 2.1.10, GTR substitution model, gamma distribution) (Price et al., 2010). The output phylogenetic tree was integrated as backbone phylogeny into PICRUSt2 to predict RRNs for all ASVs in our sample based on curated RRN values from rrnDB as reference database. The average phylogenetic distances to the closest relatives in the PICRUSt2 default reference database ranged from 0.011-0.128 for the internal reference database integrated into the PICRUSt2 software and from 0.025-0.179 for the rrnDB (abundance weighted NSTI, Table S6).
Community weighted means (CWMs) of genomic trait data in each sample were computed by multiplying the predicted trait values for each ASV with its relative abundance and adding up these weighted values in each sample. The estimation of generation time was based on the codon usage bias as detailed elsewhere (Weissman et al., 2021). Weissmann et al. suggested that only generation times up to 5 h can be reliable predicted from the codon usage bias, while the prediction of larger generation times becomes increasingly inaccurate. CWMs of generation times in our samples were in most cases ≤ 5h and reached maximally 6.2 h. We, therefore, decided that generation time estimations were sufficiently accurate.

In agreement with  earlier considerations, we classified into RRN and generation time as resilience-related traits and %TF and genome size as resistance-related traits (Beier et al., 2022). The Shannon diversity index was computed from the ASV compositional data to estimate species diversity.

### Community functional measurements

Cell growth in the continuous culture were estimated by measuring cell densities via FC as detailed elsewhere (Marie et al., 2000). Briefly, aliquots (1350 µl) were sampled using sterile syringes, fixed with glutaraldehyde (0.1% final concentration) and stored at -80°C. Samples were analyzed in the Cytoflex Flow Cytometer (Beckman Coulter, California, USA).

Functional resistance of the continuous culture was estimated weekly by measuring the below-described bulk community functional rates before and 1 hour after each induced disturbance in the disturbance treatments and simultaneously also in the controls (Table S3). Heterotroph bacterial production (BP) was assessed via $^3$H-leucine incorporation: 100 µL of a working solution containing 1 part $^3$H-leucine (125.6 Ci mmoL$^{-1}$, Perkin Elmer$^{TM}$, Massachusetts, USA) and 4 parts cold leucine were added to 1.5 mL sample water (final leucine concentration: 40 nM) and incubated in dark at 18°C. The incubations were stopped after 1.5 hours by adding 150 µL 50% trichloroacetic acid (TCA). For each reactor vessel and measure point, two technical replicates and one blanc control that was stopped by adding 50% TCA before the incubation started were performed. The incubations were then processed using the microcentrifuge method as published elsewhere (Smith and Azam, 1992), and quantified using a liquid scintillation counter (Hidex 300SL HIDEX, Turku, Finland). The theoretical conversion factor of 1.55 kg C moL$^{-1}$ was used to convert leucine incorporation rates to carbon production (Kirchman, 1993).

Respiration was quantified as the oxygen consumption estimated in 5 mL glass vials equipped with an OxoDish using a SensorDish reader (PreSens, Regensburg, Germany). The tubes were filled with sample water and closed with an air-tight lid while avoiding the enclosure of air bubbles. The filled vials were subsequently placed in an incubator at 18°C in dark. Respiration was estimated as the rate of oxygen decrease from a linear fitting from oxygen measurements taken every 3 min during 14 h. Bacterial growth efficiency (BGE in %) was estimated by dividing BP by the sum of BP and respiration, while the respiration rates were transformed to carbon consumption considering a respiratory quotient of 0.89 (Williams and Giorgio, 2005).

### Resistance index

Log-response ratios (lnR) are widely used metrics to measure the relative impact of a treatment on a given variable, independently of its metric units (Hedges et al., 1999). In this study, we applied the lnR (Hillebrand and Gurevitch, 2016) to quantify the relative change of the functional rate F before and after the pulse disturbance in the disturbed communities ($lnR_D$) and in the corresponding controls where no disturbance was introduced ($lnR_C$) at the same time intervals. For the $lnR_C$, the average of the triplicates was calculated to cover the total range of variability of the response variable.

Values of the lnR close to 0 represent a low variability of the measured functional rate, while, deviations from 0 indicate an increase (decrease) of functional rates reflected in positive (negative) lnR during the interval of consideration. We considered the difference between both ratios as resistance index for F ($RI_F$) which is similar to the effect size measurement published previously by Osenberg et al (1997). Since we were interested in a resistance metric that was independent from the direction of change, we used differently from the originally published metric the absolute difference in this study. We also did not normalize by time, as the considered time interval was equal for $lnR_D$ and $lnR_C$.

$$RI_F = -|lnR_C - lnR_D|$$

If $RI_F$ approximates zero, the temporal change of the functional response F was similar in magnitude and direction in both, the disturbed and control communities. In this case, F was only marginally impacted by the induced salinity pulse pointing to a high resistance level. The larger the deviation of $RI_F$ from zero, the lower/higher the functional resistance/sensitivity. The absolute difference between $lnR_D$ and $lnR_C$ consequently results in small value for high resistance and large values for low resistance. A multiplication of this term with -1 turned this relationship and allowed us to display the obtained RI values on a more intuitive scale where small values reflect low resistance and high values high resistance (Fig. 4).

### Statistical analysis

Statistical analyses were performed in R (R Core Team, 2018). At the end of the long-term experiment (day 41), each one replicate of the disturbance treatment under HDOM and LDOM was affected by a sudden community collapse (Fig. 4D). We have therefore considered data obtained from FC for statistical analyses only until day 40. Data obtained from the final DNA sampling event at day 41 were included in the downstream statistical analyses as it was sampled early during day 41 and most of the water volume was derived from day 39 or 40.

Compositional changes of the communities were evaluated using the Bray-Curtis distance and the phylogenetic compositional structure of the communities was assessed using a pairwise abundance weighted UniFrac metric (Lozupone and Knight, 2005). To test the effect of time, disturbance treatment, and DOM regime on the (phylogenetic) community composition, we performed Multivariate Analyses of Variance (PERMANOVA) using the function adonis2 available in the "vegan" R-package (permutations=1000, Oksanen et al., 2019).

To assess the hypothesis of the gained functional resistance under high resource conditions we performed two-ways repeated-measurements analyses of variances (rmANOVAs) on the resistance indices ($RI_F$) considering time and DOM level. We additionally fitted mixed linear models (MLM) to evaluate the direction of potentially detected trends over time, using the "nlme" R-package (Pinheiro et al., 2020). For this model, we considered time as a fixed factor and the replicates as random factors.

We furthermore performed two-way rmANOVAs to assess the effect of time and the disturbance regime on bacterial abundances, Shannon diversity, and genomic traits in each of the DOM regimes, separately. In our analysis, we aimed to test the one-tailed hypotheses predicting a simultaneous increase of resistance and resilience levels and the associated genomic trait of aquatic prokaryote communities in response to disturbances (Beier et al., 2022). More precisely we expected higher RRN, higher %TF, higher CUB and higher RRN in disturbed compared to undisturbed treatments. As a consequence of formulating one-tailed hypotheses, we performed additionally to the rmANOVAs one tail paired t-tests on the mean values of the genomic traits to test if the a priori hypothesized direction of the response values was observed. Normality and homogeneity of variance of the data were tested by the Kolmogorov-Smirnov and the Levene test, respectively. Overall no violations of the ANOVA assumptions concerning normality and homogeneity of variance were detected, except for bacterial abundance and community respiration. In these cases, a square root transformation was applied to the data to fulfill the assumptions.

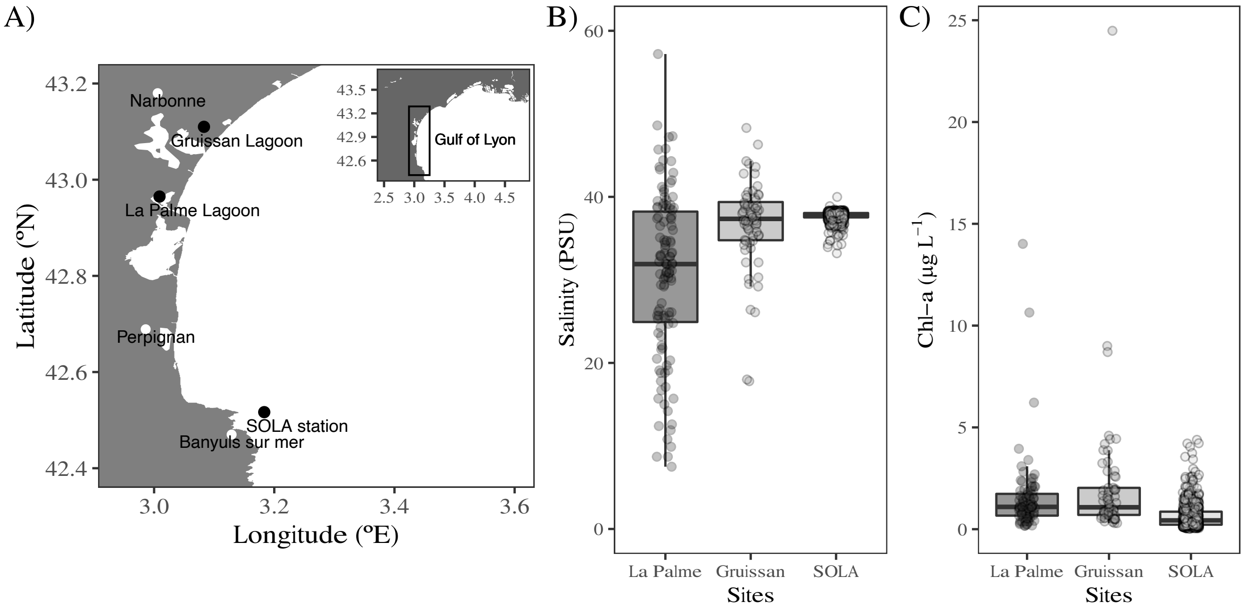


***Figure S1. Sampling locations and environmental characteristics of the sample sites, from which the starting inocula were obtained. A)*** *Geographic locations of the La Palme and Gruissan lagoons as well as the Mediterranean coastal SOLA field station.* ***B)*** *Temporal salinity (PSU) and* ***C)*** *Chlorophyll-a (µg $L^{-1}$) variability. Time series environmental data for La Palme (23/09/1989-11/09/2020, n = 206) and Gruissan (23/09/1989-08/08/2019, n = 128) were obtained from [https://wwz.ifremer.fr/surval](https://wwz.ifremer.fr/surval). Environmental data for the SOLA station was provided by the SOMLIT program ([https://www.somlit.fr/](https://www.somlit.fr/); 04/01/2005-24/11/2020, n = 1443).*

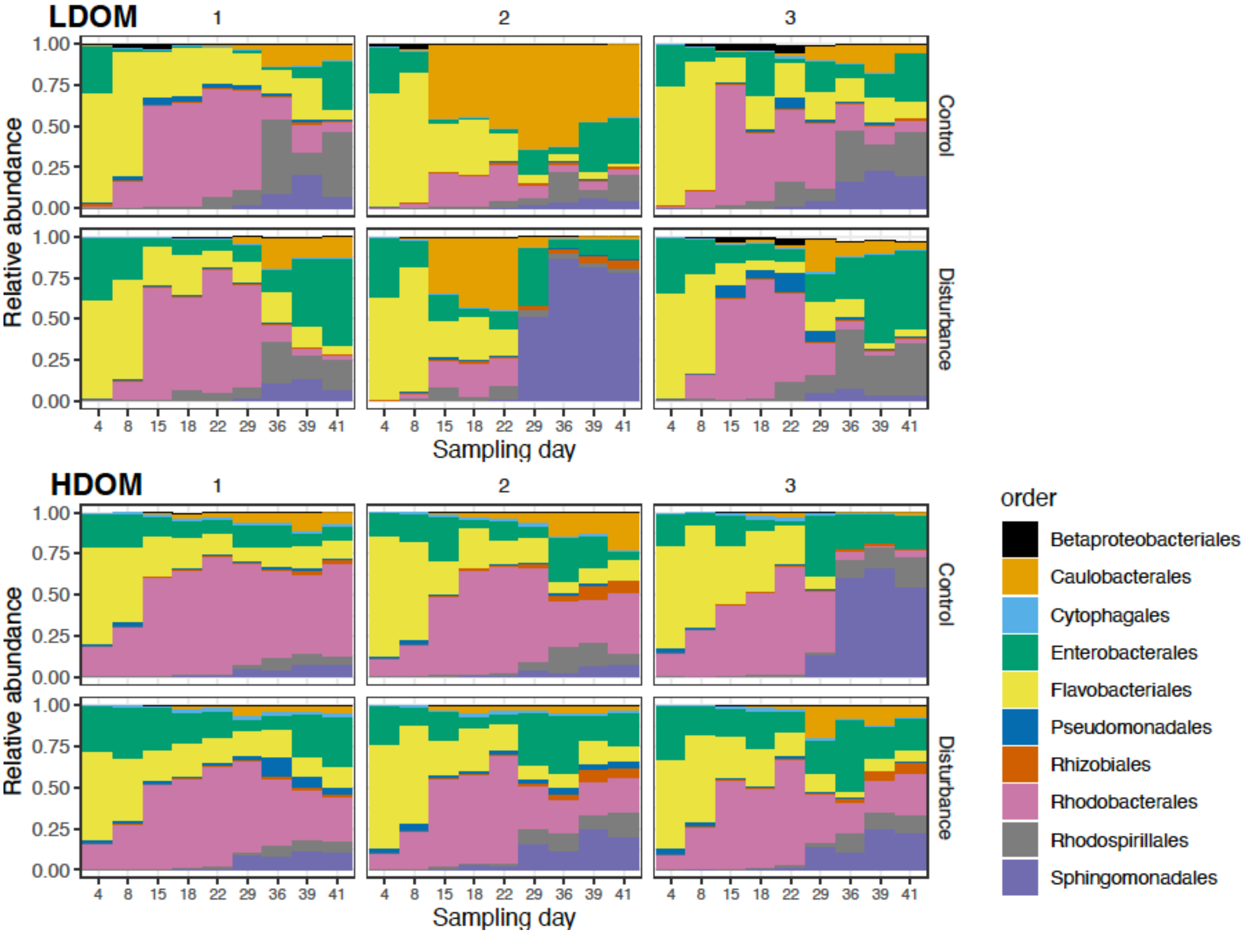


***Figure S2. Community dynamics.*** *Relative abundance of the 10 most abundant orders in the initial communities as well as in the resuscitated communities at **A)** LDOM and **B)** HDOM levels at each sampling day.*

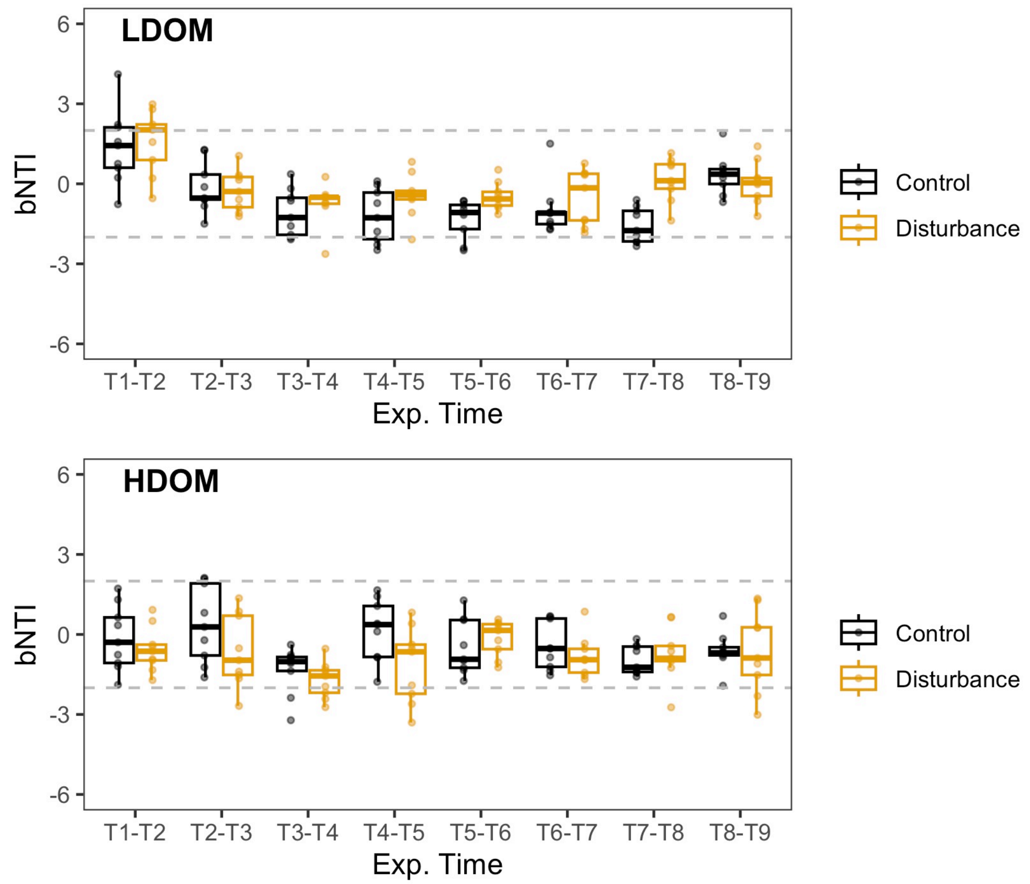


***Figure S3. Influence of deterministic versus stochastic processes on microbial community dynamics.*** *The influence of deterministic versus stochastic processes on microbial community dynamics was quantified during the course of the continuous culture experiment via null model analyses. For this purpose, we applied the bNTIs (Webb, 2000; Stegen et al., 2012) separately for disturbance and DOM regimes and over time applying a sliding window setup in the continuous cultures and using the iCAMP R-package (v1.3.4, Ning et. al., 2020).*

*The bNTI evaluates whether the phylogenetic similarity between a pair of samples is significantly lower or higher than expected by chance relative to a reference species pool. Phylogenetic similarity surpassing the theoretical expectation (bNTI >2) indicates the prevalence of variable deterministic assembly processes during community succession. Phylogenetic similarity below the theoretical expectation (bNTI< -2), indicates the prevalence of homogeneous deterministic assembly processes during community succession. bNTIs between -2 and 2 indicate that community assembly is driven by stochastic rather than a deterministic processes.*

*For our analyses bNTIs during community succession were estimated in a way that communities from each chemostat vessel were compared to the community of same vessel from the respectively following sampling time. The null models to obtain bNTIs were estimated relative to the total species pool from all incubations under the same DOM level and same disturbance regime over the incubation (=27 samples). This reference species pool was chosen because we aimed to focus on assembly mechanisms under the respectively relevant disturbance and DOM regime over time.*

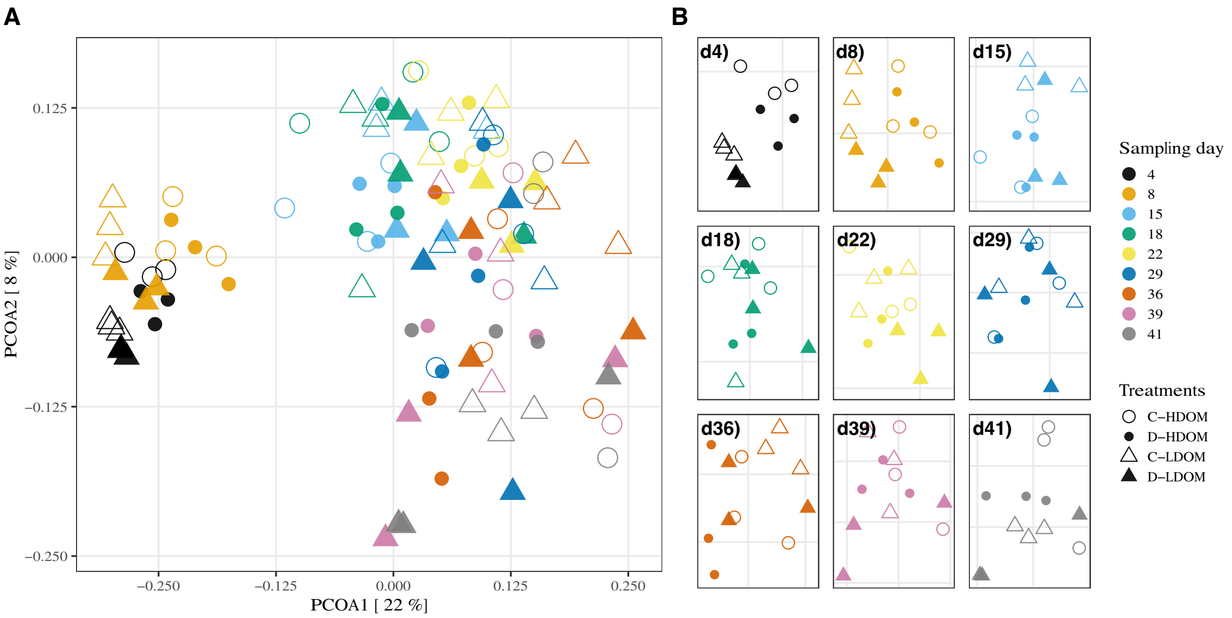


***Figure S4. Community structures.*** *A) Overview PCoA biplot including all data points (weighted Unifrac distances). B) PcoAs (weighted Unifrac distances) for individual sampling days; axes of the individual plots are differently scaled as indicated by the grid lines.*

### SI REFERENCES


Baho, D. L., Peter, H., and Tranvik, L. J. (2012). Resistance and resilience of microbial communities - temporal and spatial insurance against perturbations. *Environ. Microbiol.* 14, 2283–2292. doi: 10.1111/j.1462-2920.2012.02754.x.

Beier, S., Werner, J., Bouvier, T., Mouquet, N., and Violle, C. (2022). Trait-trait relationships and tradeoffs vary with genome size in prokaryotes. *Frontiers in Microbiology* 13.

Callahan, B. J., Mcmurdie, P. J., Rosen, M. J., Han, A. W., Johnson, A. J. A., and Holmes, S. P. (2016). DADA2 : High-resolution sample inference from Illumina amplicon data. 13. doi: 10.1038/nmeth.3869.

Douglas, G. M., Maffei, V. J., Zaneveld, J. R., Yurgel, S. N., Brown, J. R., Taylor, C. M., et al. (2020). PICRUSt2 for prediction of metagenome functions. *Nature Biotechnology* 38, 685–688. doi: 10.1038/s41587-020-0548-6.

Edgar, R. C. (2004). MUSCLE: a multiple sequence alignment method with reduced time and space complexity. *BMC Bioinformatics* 5, 1–19. doi: 10.1186/1471-2105-5-113.

Eguchi, M., Nishikawa, T., Macdonald, K., Cavicchioli, R., Gottschal, J. C., and Kjelleberg, S. (1996). Responses to Stress and Nutrient Availability by the Marine Ultramicrobacterium Sphingomonas sp. Strain RB2256. *Appl. Environ. Microbiol.* 62, 1287.

Hedges, L. V., Gurevitch, J., and Curtis, P. S. (1999). The Meta-Analysis of Response Ratios in Experimental Ecology. *Ecology* 80, 1150–1156. doi: 10.2307/177062.

Hillebrand, H., and Gurevitch, J. (2016). "Meta-Analysis and Systematic Reviews in Ecology," in *eLS*, ed.John Wiley & Sons Ltd (Chichester, UK: John Wiley & Sons, Ltd), 1–11. doi: 10.1002/9780470015902.a0003272.pub2.

Kirchman, D. (1993). *Handbook of Methods in Aquatic Microbial Ecology*.

Landa, M., Cottrell, M. T., Kirchman, D. L., Kaiser, K., Medeiros, P. M., Tremblay, L., et al. (2014). Phylogenetic and structural response of heterotrophic bacteria to dissolved organic matter of different chemical composition in a continuous culture study. *Environ Microbiol* 16, 1668–1681. doi: 10.1111/1462-2920.12242.

Lozupone, C., and Knight, J. R. (2005). UniFrac: a new phylogenetic method for comparing microbial communities. *Applied and Environmental Microbiology* 71, 8228–8235.

Marie, D., Simon, N., Guillou, L., Partensky, F., and Vaulot, D. (2000). "Flow Cytometry Analysis of Marine Picoplankton," in *In Living Color: Protocols in Flow Cytometry and Cell Sorting* Springer Lab Manuals., eds. R. A. Diamond and S. Demaggio (Berlin, Heidelberg: Springer), 421–454. doi: 10.1007/978-3-642-57049-0_34.

Ning, D., Yuan, M., Wu, L., Zhang, Y., Guo, X., Zhou, X., et al. (2020). A quantitative framework reveals ecological drivers of grassland microbial community assembly in response to warming. *Nature Communications* 11, 4717. doi: 10.1038/s41467-020-18560-z.

Oksanen, J., Blanchet, F. G., Friendly, M., Kindt, R., Legendre, P., McGlinn, D., et al. (2019). *vegan: Community Ecology Package*. R package version 2.5-6. Available at: https://CRAN.R-project.org/package=vegan.

Osenberg, C. W., Sarnelle, O., and Cooper, S. D. (1997). Effect Size in Ecological Experiments: The Application of Biological Models in Meta-Analysis. *The American Naturalist* 150, 798–812. doi: 10.1086/286095.

Parada, A. E., Needham, D. M., and Fuhrman, J. A. (2016). Every base matters : assessing small subunit rRNA primers for marine microbiomes with mock communities , time series and global field samples. 18, 1403–1414. doi: 10.1111/1462-2920.13023.

Parks, D. H., Chuvochina, M., Waite, D. W., Rinke, C., Skarshewski, A., Chaumeil, P.-A., et al. (2018). A standardized bacterial taxonomy based on genome phylogeny substantially revises the tree of life. *Nature Biotechnology* 36, 996–1004. doi: 10.1038/nbt.4229.

Pinheiro, J., Bates, D., DebRoy, S., Sarkar, D., and R Core Team (2020). {nlme}: Linear and Nonlinear Mixed Effects Models. Available at: https://CRAN.R-project.org/package=nlme.

Price, M. N., Dehal, P. S., and Arkin, A. P. (2010). FastTree 2-Approximately Maximum-Likelihood Trees for Large Alignments. *PLoS One* 5. doi: 10.1371/journal.pone.0009490.

R Core Team (2018). R: A Language and Environment for Statistical Computing. Available at: https://www.R-project.org/.

Rain-Franco, A., de Moraes, G. P., and Beier, S. (2021). Cryopreservation and Resuscitation of Natural Aquatic Prokaryotic Communities. *Frontiers in Microbiology* 11, 3633. doi: 10.3389/fmicb.2020.597653.

Smith, D., and Azam, F. (1992). A simple economical method for measuring bacterial protein synthesis rates in seawater using 3H-Leucine. *Marine Microbial Food Webs* 6, 107–114.

Stegen, J. C., Lin, X., Konopka, A. E., and Fredrickson, J. K. (2012). Stochastic and deterministic assembly processes in subsurface microbial communities. *The ISME Journal* 6, 1653–1664. doi: 10.1038/ismej.2012.22.

Stoddard, S. F., Smith, B. J., Hein, R., Roller, B. R. K., and Schmidt, T. M. (2015). rrnDB: improved tools for interpreting rRNA gene abundance in bacteria and archaea and a new foundation for future development. *Nucleic Acids Res.* 43, D593–D598. doi: 10.1093/nar/gku1201.

Webb, C. O. (2000). Exploring the Phylogenetic Structure of Ecological Communities: An Example for Rain Forest Trees. *The American Naturalist* 156, 145–155. doi: 10.1086/303378.

Weissman, J. L., Hou, S., and Fuhrman, J. A. (2021). Estimating maximal microbial growth rates from cultures, metagenomes, and single cells via codon usage patterns. *PNAS* 118. doi: 10.1073/pnas.2016810118.

Williams, P. J. le B., and Giorgio, P. A. del (2005). *Respiration in aquatic ecosystems: history and background*. Oxford University Press Available at: https://oxford.universitypressscholarship.com/view/10.1093/acprof:oso/9780198527084.001.0001/acprof-9780198527084-chapter-1 [Accessed June 8, 2021].
